# A simple test identifies selection on complex traits in breeding and experimentally-evolved populations

**DOI:** 10.1101/238295

**Authors:** Tim Beissinger, Jochen Kruppa, David Cavero, Ngoc-Thuy Ha, Malena Erbe, Henner Simianer

## Abstract

Important traits in agricultural, natural, and human populations are increasingly being shown to be under the control of many genes that individually contribute only a small proportion of genetic variation. However, the majority of modern tools in quantitative and population genetics, including genome wide association studies and selection mapping protocols, are designed to identify individual genes with large effects. We have developed an approach to identify traits that have been under selection and are controlled by large numbers of loci. In contrast to existing methods, our technique utilizes additive effects estimates from all available markers, and relates these estimates to allele frequency change over time. Using this information, we generate a composite statistic, denoted *Ĝ*, which can be used to test for significant evidence of selection on a trait. Our test requires pre- and post-selection genotypic data but only a single time point with phenotypic information. Simulations demonstrate that *Ĝ* is powerful for identifying selection, particularly in situations where the trait being tested is controlled by many genes, which is precisely the scenario where classical approaches for selection mapping are least powerful. We apply this test to breeding populations of maize and chickens, where we demonstrate the successful identification of selection on traits that are documented to have been under selection.

## Introduction

Quantitative traits encompass an inexhaustible number of phenotypes that vary in populations, from characters such as height (Yang *et al.* 2010) to weight (Barsh *et al.* 2000) to disease resistance (Poland *et al.* 2009). These types of traits are so essential for agriculture and human health that the entire field of quantitative genetics revolves around their study (Plomin *et al.* 2009; Wallace *et al.* 2014). However, the nature of quantitative traits makes it difficult to study their genetic basis; for nearly a century, scientists have modeled quantitative traits by assuming that their underlying control involves many loci each contributing a very small proportion to genetic variance (Fisher 1918), the so-called ‘infinitesimal model’. Therefore, conducting studies with enough power to identify a substantial proportion of the loci that contribute to a quantitative trait requires a massive sample size, imposing financial and logistical barriers. However, this model of quantitative trait variation does an excellent job when predicting important characteristics such as response to selection (Visscher *et al.* 2008). For instance, genomic prediction methodologies (Meuwissen *et al.* 2001) allow the breeding value and/or phenotype of individuals to be predicted with remarkable precision from genomic information alone.

The models of quantitative genetics have had a less dramatic impact on studies of evolutionary adaptation, where genomes are often scanned to identify adaptive loci with large effects (Akey 2009). Positive selection on such loci leaves behind pronounced signatures, deemed “selective sweeps”. There is an abundance of evidence for such sweeps in humans (Sabeti *et al.* 2007), natural populations (Schweizer *et al.* 2016), livestock (Qanbari and Simianer 2014), and crops (Hufford *et al.* 2012; Qanbari and Simianer 2014; Schweizer *et al.* 2016). However, alternative forms of selection, including purifying selection against new mutations (Lawrie *et al.* 2013), selection on standing variation (Garud *et al.* 2015), or selection on many loci of small effect (Turchin *et al.* 2012), rarely leave these discernible signatures at individual loci. Evidence of these forms of selection can be difficult to identify. When they can be found, it is often through the pooling of weak evidence at individual loci into a stronger signal across a class of loci. For example, Beissinger et al (2016) demonstrated the importance of purifying selection during maize evolution by combining evidence from all maize genes. An approach implemented by Berg and Coop (2014) tests for evidence of selection on a quantitative trait by evaluating allele frequencies at all loci that have previously been implicated by genome-wide association studies (GWAS) as putatively associated with that trait. This approach has since been used to test for selection on multiple human traits, including height (Mathieson *et al.* 2015) and telomere length (Hansen *et al.* 2016).

In studies of model organisms or agricultural species, large collections of previously identified “GWAS hits” are not as abundant as in humans, on which the Berg and Coop (2014) method depends. This is due in part to the more modest sample sizes that tend to be used in experimental settings compared to clinical studies, often combined in large-scale meta-analyses (Evangelou and Ioannidis 2013). Conversely, genotypic data across at least two time points are often readily available for model and agricultural species. Due to improving technologies for sequencing ancient DNA (Mathieson *et al.* 2017; Berg *et al.* 2017), and/or by leveraging populations that have benefitted from excellent historical record-keeping (Kong *et al.* 2017), genetic data with a temporal component is increasingly available in humans. We have developed a test for selection on complex traits that leverages such genotype-over-time data. Our test depends on the relationship between the change in allele frequency between two generations and the estimated additive effect of the same allele, computed for every genotyped locus. We use these values to compute an estimate of the direction of genetic gain, which can be shown to be additive across all loci considered. Our estimate lends itself to a simple permutation-based test for significance that avoids many of the demographic history and population structure related caveats that complicate determining significance when testing for selection (de Villemereuil *et al.* 2014). The method utilizes additive effects estimates for each locus calculated simultaneously, using shrinkage-based methods that have been honed over the past 15 years for the purpose of genomic selection and prediction (Campos *et al.* 2013). Therefore, this test can be considered analogous to reverse genomic selection; rather than using predictions of breeding value to drive selection and hence future changes in allele frequency, we use the same data coupled with knowledge of past changes in allele frequency to make inferences regarding which traits were effectively under selection in the past. Interestingly, we find by simulation that this approach is most powerful for identifying selection on traits controlled by many loci of small effect, which is exactly the situation where other tests for selection and/or association are least powerful.

Herein, we first motivate and describe our test for selection on complex traits, which we call *Ĝ*. Then, we perform simulations demonstrating the validity of the method and explore the situations where it is most and least powerful. Finally, we apply the method to breeding populations of maize and chicken. In both of these experimental situations, we successfully identify the traits that are known to have been selected. Collectively, our results demonstrate that this approach may be leveraged to identify novel traits or component-traits that may be used to inform future breeding decisions and/or for enhanced historical, ecological, and basic scientific understanding. Software for implementing this test can be found in the accompanying Github repository, github.com/timbeissinger/ComplexSelection.

## Results

### Theoretical Motivation

Assume that a trait is fully controlled by additive di-allelic loci *j =* 1,… *m.* Then the genotypic value, *a_j_,* of an allele at locus *j,* is equal to its gene substitution effect, *α_j_.* Based on this equivalency, the mean phenotypic effect, *M_j_,* attributable to the locus is given by *M_j_ = α_j_(2p_j_-1),* where *p_j_* is the frequency of the reference allele at this locus. It follows that the change in the population mean resulting from selection on this locus, what we may consider the locus-specific response to selection, is given by

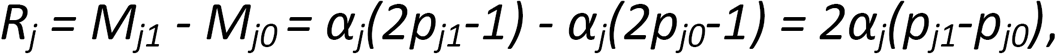

where *p_j0_* is the allele frequency before selection and *p_j1_* is the allele frequency after selection. Define *Δ_j_ = (p_j1_-p_j0_),* leading to *R_j_* = 2 *Δ_j_α_j_.* Based on our earlier assumption of complete additivity, summing over all m loci provides a genome-wide estimate of the response to selection (Falconer and Mackay 1996):

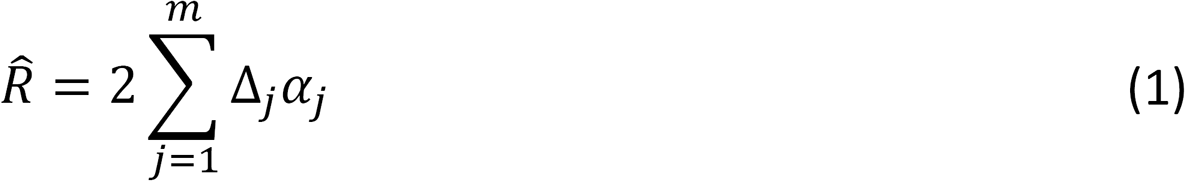

Strictly speaking, since relative effect sizes may change each generation with changing allele frequencies throughout the genome, (1) is applicable for a single generation. However, under the assumption of many loci affecting a trait, (1) may approximately apply for many generations of selection. This estimate of selection response also naturally arises from the logic of random regression BLUP (RRBLUP) (Meuwissen *et al.* 2001). Here, a model is used

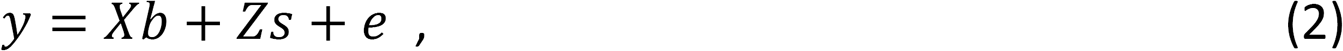

where ***y*** is a vector of length *n* containing phenotypes for a specific trait, *b* are fixed effects, 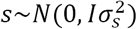 is the vector of length *m* containing additive SNP effects at *m* loci; 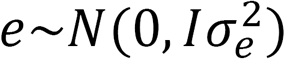 is the vector of random residual terms, and 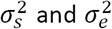 are the corresponding variance components. *X* and *Z* are incidence matrices linking observations in *y* to the respective levels of fixed effects in *b* and random SNP effects in *s.* In more detail, *Z* is an *n* × *m* matrix where element *z_ij_* contains the genotype of individual *i* at SNP locus *j.* Since such models are invariant with respect to linear transformations of the allele coding (Strandén *et al.* 2011), we may use the notation 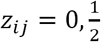, *or* 1, standing for zero, one, or two copies of the reference allele. Note that with this coding, *S_j_* is equivalent to 2*α_j_* in the coding above, since it reflects the contrast between the two homozygous genotypes at locus *j.* Due to the equivalence of genomic BLUP (GBLUP; VanRaden 2008) and RRBLUP (Endelman 2011), it is possible to calculate genomic breeding values of the genotyped individual as *û = Zŝ,* where *ŝ* are the solutions for the SNP effects obtained using RRBLUP with model (2).

Assume now further that individuals in the vector *y* can be assigned to *g* discrete generations and that the individuals of the oldest generation come first and the individuals of the last generation come latest. We then can define a *g × n* matrix

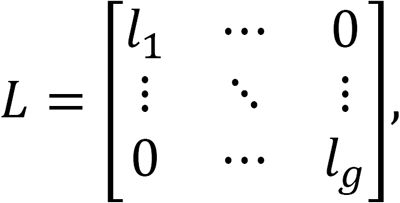

where *l_p_* is a row vector of length *n_p_,* which is the number of individuals in generation *p*, of which all elements are *1/n_p_*. With this, a vector *ū* of length *g* reflecting average breeding values per generation can be calculated as *ū = Lû,* and estimated selection response results as 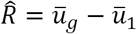. Now, *ū = Lû = LZŝ,* where *LZ* is a *g × m* matrix in which element *p, j* reflects the average allele frequency of the reference allele at SNP *j* in generation *p*. The allele frequency change between generation 1 and generation g can be obtained as a linear contrast between the first and the last row of this matrix as *Δ = k’LZ* where *k* is a vector of length g with *k_1_ = -1, k_g_ = 1*, and all other elements are zero. Finally, the selection response can be written as 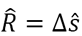 which is identical to equation (1), given that *s* is equivalent to *2α.*

Furthermore, theory suggests that under the assumption that selection intensity is equal for all loci across the genome, the change of allele frequency *Δ_j_* should be approximately proportional to the allele effect *α_j_*, such that for a trait under selection a non-zero correlation between allele frequency change and the additive effect of alleles on that trait is expected (Wright 1937). Alternatively stated, (1) emphasizes the temporal component of the Breeder’s Equation, *R = h^2^S,* where *h^2^* is the narrow-sense heritability of a trait and *S* is the selection differential. Given a population of individuals with two time-points of genotypic data, it is simple to compute *Δ_j_* for every genotyped locus. Furthermore, the shrinkage methods of genomic prediction (Campos *et al.* 2013), including Ridge Regression (Endelman 2011) and GBLUP (VanRaden 2008) allow additive effects, *α_j_,* to be approximated for every genotyped position. For this, a set of individuals genotyped and phenotyped in at least one generation is needed.

A notable benefit of the estimator in (1) is that by leveraging pre- and post-selection data from genotypes rather than from phenotypes, it only requires one generation of phenotyping. Additionally, this suggests that if we consider *R* a random variable, then given the distribution of *R* in a scenario without selection, a test of whether or not 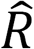 is different from zero may be performed. Since 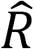 is the genomic response to selection, this is equivalent to testing whether or not a trait has been under selection during the timeframe under study.

### Test Statistic and Significance Testing

We implemented a permutation-based strategy to test whether or not 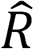 is significantly different from zero. Genetic drift and selection jointly determine changes in allele frequency, *Δ_j_*, but without selection these changes in frequency should not be related to effect size or direction. The reverse is also true; effect sizes, *α_j_*, are estimated based on a genomic prediction model applied to phenotypes measured in a single panel of individuals, and therefore they are not correlated with changes in allele frequency. While a correlation between minor allele frequency and the magnitude of SNP effects is possible due to estimation error during genomic prediction, without ongoing selection allele frequency should not correlate with the direction of SNP effects. This suggests that a null distribution for 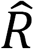 in a no-selection scenario may be generated via a permutation approach. Assuming no linkage disequilibrium (LD) between markers, a simple shuffling of *Δ_j_* and *α_j_* can be implemented to generate the desired null distribution. However, LD between markers compromises the applicability of this simplified approach for most populations—such an approach overestimates the sample size of the permutation test by treating each marker as an independent observation, while in reality any level of LD between markers leads to fewer independent observations than markers. Therefore, we have employed a semi-parametric approach that scales the variance of the permutation test statistic according to the realized extent of LD to alleviate this discrepancy.

Let 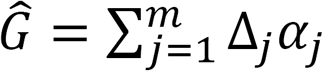, which is proportional to 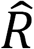 as defined in (1). This value, colloquially *“G-hat”,* serves as our test statistic. The summation is over all m genotyped markers, and effect sizes are estimated based on genomic prediction using available phenotypes with corresponding genotypes from any generation. Often, phenotypes from the most recent generation will be the most readily available, but individuals with phenotypes scored in any generation may suffice. To test whether or not the observed value of *Ĝ* can be significantly attributed to selection, define **p** to be a vector of length m that is a permutation of the vector **J** = [1,..,m]. A permuted value of *Ĝ* may be obtained via 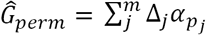. Because *Δ_j_* and *α_pj_.* are no longer indexed to the same locus, *Ĝ_perm_* does not reflect selection, but instead captures genetic drift over time (*Δ_j_* terms) as well as the genetic architecture of the underlying trait (*α_j_* terms). Generating repeated values of *Ĝ_perm_* through repeated permutations of **J** therefore generates a null distribution for *Ĝ* which assumes no selection and complete linkage equilibrium.

The Central Limit Theorem dictates that realizations of *Ĝ_perm_* are normally distributed with approximate mean 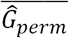 and standard deviation *SE(Ĝ_perm_)*. Therefore, σ, the underlying standard error of a single-locus estimate for *Ĝ_perm_* is given by 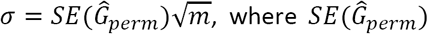 is the observed standard error of *Ĝ_perm_*. Consider the quantity *m_ind_,* representing the effective number of independent loci. If the standard deviation of *Ĝ_perm_* were calculated using *m_ind_* independent markers, its expectation would be 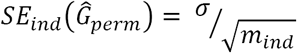.Plugging in the estimate for *σ* obtained above, *SE_ind_(Ĝ_perm_)* becomes 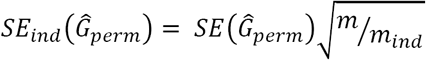.

In practice, the above implies that to test for selection, 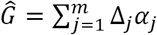 may be calculated from data, and then a permuted null distribution for *Ĝ* that assumes linkage equilibrium can be generated. This permutation distribution may then be approximated with a normal distribution, whose variance can be scaled according to the effective number of independent markers, *m_ind_*. We show in the following section that *m_ind_* can be efficiently estimated based on LD-decay. Ultimately, significance may be evaluated by comparing *Ĝ* to a normal distribution with mean 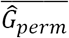 and standard deviation 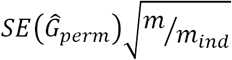.

### Simulations

We conducted a series of simulations to evaluate the power of the *Ĝ* statistic for identifying selection on complex traits. Genotypic data were simulated with the software program QMSim (Sargolzaei and Schenkel 2009). An overview of our simulation strategy at the most general level is that we simulated selection in a generic species with 1,000 QTL dispersed along ten 100 cM chromosomes, with a total of 100,000 equally-spaced markers (10,000 per chromosome). In the first step of each simulation, the total population was established based on 10,000 individuals randomly mating for 5,000 generations. Then, 500 males and 500 females were randomly chosen to establish a base population that would undergo selection for 20 generations. Each generation, 1,000 individuals (500 males and 500 females) were permitted to mate out of a population of 5,000, providing a selection proportion of 20%. For each simulation, heritability was set to 0.5. This general scheme encapsulates characteristics of most plant and animal breeding populations, including the large number of progeny typical of plants and the truncation selection protocol often associated with animal breeding and/or selection in the wild. Additional details regarding the simulated population are included in Supplemental Table 1. In the following subsections, we describe how varying parameters from the generic scenario described here affected the power of *Ĝ* to identify selection. All simulation scripts can be found at github.com/timbeissinger/ComplexSelection.

#### Number of QTL

We simulated variable numbers of additive QTL controlling traits, from 10, representing a simple trait controlled by large-effect QTL, to 10,000, representing a highly quantitative trait controlled nearly infinitesimally. QTLs were evenly spaced along each chromosome and QTLs themselves were not included in the marker set for analysis. One hundred simulations were performed for each level of trait complexity. First, we used these simulations to establish the appropriate number of independent markers, *m_ind_* as defined above, for this test. We calculated how distant two markers must be to have an expected LD level of *R^2^ ≤* 0.03. Then we counted the total number of blocks of this size genome-wide. The 0.03 level was established by performing a grid-search of potential values and tuning the false positive rate (Supplemental Figure 1). An LD cutoff that is too high leads to a high false-positive rate, while one that is too low weakens the power of the test. For populations similar to those discussed herein, we observe that requiring *R^2^ ≤* 0.03 will be appropriate.

When we tested for selection in our simulated data, we observed a direct relationship between the number of QTL controlling a trait and the power of *Ĝ* to identify selection on that trait. *Ĝ* powerfully identifies selection on highly polygenic traits, but is not powerful for identifying selection on traits controlled by a small number of QTLs. Analyses of the same simulations using *F_ST_*-based selection mapping, which involves mapping loci that have been previously subjected to selection (Wisser *et al.* 2008), showed that traits controlled by a small number of QTLs can be mapped using traditional selection mapping approaches. However, as traits become increasingly polygenic, our simulations demonstrate that the ability to map individual selected genes diminishes (Figure 1). These findings demonstrate how *Ĝ* and traditional selection mapping can be complementary depending on the underlying genetic architecture of a trait. Table 1 depicts detection and false positive rates for *Ĝ* and *F_ST_-*based mapping under different genetic architectures.

**Table 1:** Detection and false positive rates for Ĝ and selection mapping. One *Ĝ* test is conducted per simulation, so true and false positive rates are shown. For selection mapping, one test is conducted per marker in each simulation, so the mean number of markers that were declared true and false positives is shown. A marker was declared a false positive in selection mapping if it exceeded a 5% simulation-based experiment-wide significance threshold but was not within a .1 cM region around a simulated QTL. Note that there are no selection mapping false positives in the 10,000 QTL simulation because every marker was within 0.1 cM of a simulated QTL.

| Genetic Architecture | 10 QTL | 50 QTL | 100 QTL | 1,000 QTL | 10,000 QTL |
| --- | --- | --- | --- | --- | --- |
| **$\hat{G}$** |  |  |  |  |  |
| True positive rate | 0.04 | 0.54 | 0.94 | 1.0 | 1.0 |
| False positive rate | 0.03 | 0.03 | 0.02 | 0.03 | 0.04 |
| **$F_{ST}$-based Selection Mapping** |  |  |  |  |  |
| Mean # true positives (rate) | 5.6 (56%) | 22 (44%) | 39 (39%) | 187 (18.7%) | 1,676 (16.8%) |
| Mean # false positives | 52 | 280 | 715 | 1,745 | - |

**Figure 1:**
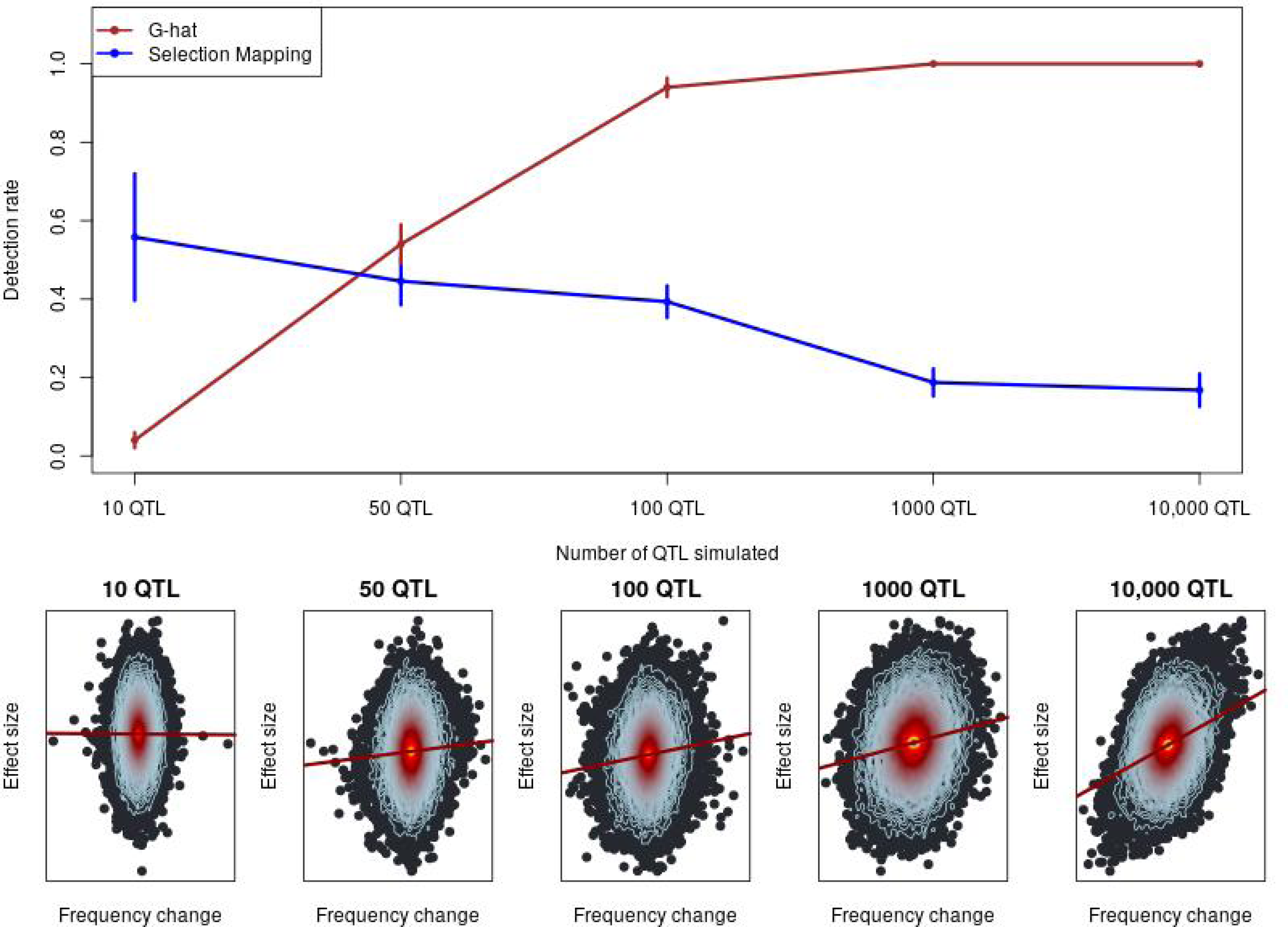
The power of Ĝ to identify selection. Top: The detection rate of Ĝ compared to Fst-based selection mapping. Vertical lines indicate one standard deviation. Standard deviations for selection mapping were estimated empirically. Standard deviations for Ĝ were estimated based on the binomial distribution. Bottom: Exemplary heat plots depicting individual-SNP allelic effect estimates linearly regressed on allele frequency change over time. Each point represents a SNP, while the contour lines indicate the density of SNPs. From the regression line, observe that a stronger relationship between frequency change and effect size corresponds to increasing polygenicity.

#### Number of generations

Simulations showed an interesting relationship between the number of generations of selection and the power of *Ĝ*. We observed a definite sweet-spot from ~10 to just under 50 generations for which *Ĝ* was most powerful. Conversely, if selection took place for 100 generations or only for a single generation, *Ĝ* became dramatically less powerful (Table 2). We suspect that two forces interact to reduce the power of *Ĝ* in the case of a large number of generations of selection. First, over the course of many generations, our simulated populations became highly inbred, which notably increased LD and therefore reduced *M_ind_.* Since *Ĝ* is summed over markers and then scaled by *M_ind_,* this substantially reduces power. Secondly, our simulations involved a predetermined number of QTL with fixed effects at the onset of selection, but as selection persisted these QTL could be lost to fixation, or as allele frequencies change, their effects could decrease (Sargolzaei and Schenkel 2009). Since we estimated SNP effects based on phenotypes in the final generation (but see the section on phenotyping generation, below), power could be reduced by the fixation of a lost QTL that previously had an effect. Although these issues weakened *Ĝ* in our simulations, it is unclear whether or not they would have the same impact in a real application, and it is likely unlikely that the powerful sweet-spot would be the same. Regarding the weak power of *GĜ* to identify selection after only one generation, this is not unexpected, since for quantitative traits a single generation is rarely long-enough to appreciably shift allele frequencies.

**Table 2:** Power of *Ĝ* as simulation parameters vary. Aside from whichever parameter was being explored, simulations assumed 20 generations of selection with a selection intensity of 0.2, a genetic architecture of 1,000 QTL, a selection population consisting of 500 males and 500 females, and the additional parameters of our “generalize” selection scenario are given in Supplementary Table 1.

| **Parameter Varied** | **Tested Values** |  |  |  |  |
| --- | --- | --- | --- | --- | --- |
| **No. individuals phenotyped** | **1,000** | **500** | **250** | **100** | **50** |
| **Power** | 1 | 0.99 | 0.83 | 0.4 | 0.21 |
| **Selection intensity** | **1%** | **5%** | **20%** | **50%** | **-** |
| **Power** | 0.95 | 0.99 | 1.0 | 1.0 | - |
| **No. Gens. of selection** | **100** | **50** | **20** | **10** | **1** |
| **Power** | 0 | 0.81 | 1.0 | 1.0 | 0.18 |
| **Phenotyping generation** | **20** | **10** | **0** | **-** | **-** |
| **Power** | 1 | 1 | 0.86 | - | - |
| **No. Gens. post-selection** | **5** | **20** | **50** | **100** | **-** |
| **Power** | 1 | 1 | 0.26 | 0 | - |

We also investigated how the power of *Ĝ* is affected by temporary selection. Specifically, we simulated 20 generations of selection followed by different numbers of generations without selection. We observe that *Ĝ* remains powerful for at least 20 generations post-selection, but after 100 generations without selection, the ability of *Ĝ* to identify selection is lost. Like above, this loss of power can likely be attributed to inbreeding and the fixation of QTL.

#### Phenotyping generation

In practical applications, we predict that phenotypes will typically be more readily available from later generations of selection than early generations. However, since this generalization will not always apply, we explored how the power of *Ĝ* is affected by the generation in which individuals are phenotyped. We observed the highest power when phenotypes were scored in recent time-points or midway through selection, but power was still high (0.86) when phenotypes were scored in generation 0, at the onset of selection (Table 2). As discussed above in the section on the number of generations of selection, changing QTL effects as allele frequencies change during evolution are likely to explain this drop in power. We explored whether or not the generation of phenotyping can lead to bias by evaluating the false positive rate for simulations where phenotypes were scored at different time-points, out of 20 generations of selection. False positive rates were respectively 0.02, 0.08, and 0.0, when phenotyping occurred in generation 20, 10, and 0.

#### Intensity of selection

The intensity of selection, or the proportion of individuals that reproduce each generation, directly impacts the efficacy of a selection regime. Therefore, we explored the ability of *Ĝ* to identify selection across several selection intensities representing realistic values observed in experimental and agricultural selection programs (Table 2). To achieve this, in our simulations we varied the total number of progeny each generation rather than altering the total number of individuals reproducing, as a reduced number of individuals would rapidly lead to high levels of inbreeding. For intermediate to strong selection intensities, from 50% to 5% of individuals reproducing, we observed that *Ĝ* was highly effective for identifying selection, with power at or near 1.0. Only in the case of very strong selection, when 1% of individuals reproduced each generation, did we observe a minor reduction in the power of *Ĝ.* Despite our attempts to minimize inbreeding in these simulations, in the case of 1% selection intensity inbreeding was likely still generated via a large number of progeny originating from the same combination of superior parents. We suspect this is what resulted in the reduction in power.

#### Sample size

Since the accuracy of estimated marker effects depends on sample size, we explored the impact that the number of phenotyped individuals has on the power of *Ĝ*. Unsurprisingly, as sample size decreases so does the power of *Ĝ* to identify selection (Table 2). However, it is notable that even with sample sizes as small as 250 individuals, power remains above 0.8. Even with only 50 phenotyped individuals, selection can be identified in one out of five scenarios. Together, these observations emphasize that the power of *Ĝ* comes from its accumulation of information across markers rather than from a small number of highly-informative markers.

### Selection on maize silage traits

We re-analyzed data from a previous study that tested for selection in a decades-long breeding program for maize silage quality (Lorenz *et al.* 2015). Very briefly, a selection index comprised of experimentally-measured traits related to silage quality was used to perform reciprocal recurrent selection for breeding improved maize. Traits comprising the index included acid detergent fiber (ADF), protein content, starch content, *in-vitro* digestibility, and yield (www.cornbreeding.wisc.edu). In total, 648 individuals from various stages of selection were genotyped. Between 240 and 300 of these individuals were also phenotyped, depending on the trait. Selection mapping was previously performed utilizing simulations of drift to scan for selection, but the analysis did not identify any loci that showed significant evidence of selection. This is in spite of quantifiable improvement of the population and demonstrated heritability of the index-composing traits (Lorenz *et al.* 2015). We re-analyzed the same data to evaluate evidence for polygenic selection on the measured traits, which included NDF, *in-vitro* digestibility, crude protein content, starch content, yield, and dry matter. After filtering for quality, but not minor allele frequency, these data consisted of 10,023 polymorphic markers. Genomic prediction for these traits was generally effective (Supplemental Figure 2). Due to the relatively small population size and recurrent selection breeding scheme, we expect slow LD decay and therefore for most of the genome to be represented with this marker set. Further analysis of LD to determine the value of *m_ind_* to utilize in our test for selection confirms this (Supplemental Figure 3).

Figure 2 depicts the maize patterns of selection that were observed in our analysis. In these plots, the histogram shows the null distribution of *Ĝ* that was observed from a permutation test, while the vertical line depicts the observed value of *Ĝ* when applied to the experimental data. We observed that with the exception of protein, for the traits where we had an *a priori* expectation of selection, we not only identified that selection did occur, but we correctly estimated the direction of selection (positive or negative) from the data. One of the traits measured was silage dry matter (DM), which was not a part of the selection index. We did not identify evidence of selection on DM, as was expected. To ensure that the existence of a single individual with a high breeding value does not lead to spurious false positives, we re-analyzed the maize data after removing all SNPs with minor allele frequency less than 0.05. This did not lead to any appreciable change in the results (Supplemental Figure 6).

**Figure 2:**
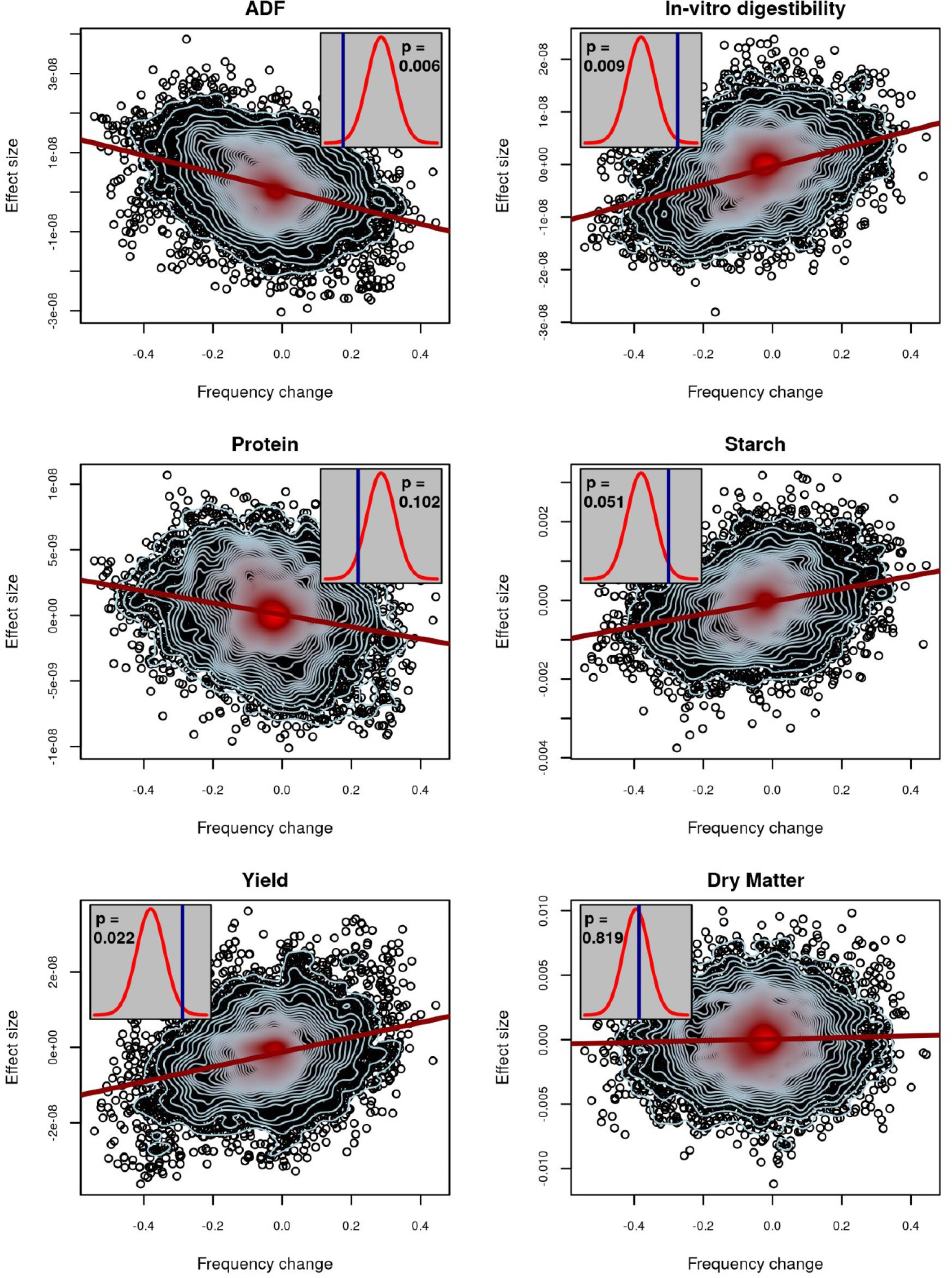
Evidence of selection for maize silage traits. For six traits, the relationship between estimated allelic effects at individual SNPs and the change in allele frequency over generations is plotted. The red line is a regression of effect size on allele frequency change. Contour lines indicate the density of points, with blue contours indicating fewer points than red. Inset plots depict observed values of Ĝ (blue lines) and their statistical significance based on a comparison to permuted null distributions (red densities) for no-selection scenarios. An exact two-sided p-value is given within each inset. Significant values of Ĝ above the permuted mean indicate selection operated in the positive direction, while significant values below the permutation mean indicated selection operated in the negative direction.

### Selection on chicken traits

We tested for evidence of selection in two panels of commercial lines of laying hens: one white layer (WL) and one brown layer (BL). Both closed lines have been selected over decades with a similar composite breeding goal, comprised of laying rate, body weight and feed efficiency, egg weight, and egg quality, among other objectives. The respective weights applied to the different traits varied between lines and over time. Traits analyzed included laying rate, egg weight, and breaking strength of eggs. Genotypes were available only for the post-selection population, so initial allele frequencies were inferred based on pedigree data (Gengler *et al.* 2007). *M_ind_* was determined based on separate evaluations of LD in the WL (Supplemental Figure 4) and BL (Supplemental Figure 5) populations.

Among the traits evaluated, we observed significant evidence of selection for increased laying rate in both WLs (p = 0.021) and BLs (p = 0.021). Tests were also suggestive of selection for increased eggshell breaking strength in WLs (p < 0.1; one-sided p < 0.05), while there was no evidence of directed selection for egg weight (Figure 3). To verify that these results were not driven by a small number of SNPs with high estimated effect sizes, we repeated the analysis with the 10 largest effect-size SNPs removed and saw virtually identical results (Supplemental Figure 7). The result for egg weight can be seen as a ‘negative control’ since for this trait an optimum value is already achieved and maintained by stabilizing selection. The fact that we were not able to detect significant evidence of selection in a trait such as eggshell breaking strength in both lines (although a tendency can be observed) may be due to the fact that improving those traits is part of a complex multi-objective breeding program, or simply that our test was underpowered for these traits. The unavailability of experimentally-estimated initial frequencies and our alternative use of pedigree-inferred initial allele frequencies likely weakened the power of the test as compared to the more complete data available for maize and in the simulations.

**Figure 3:**
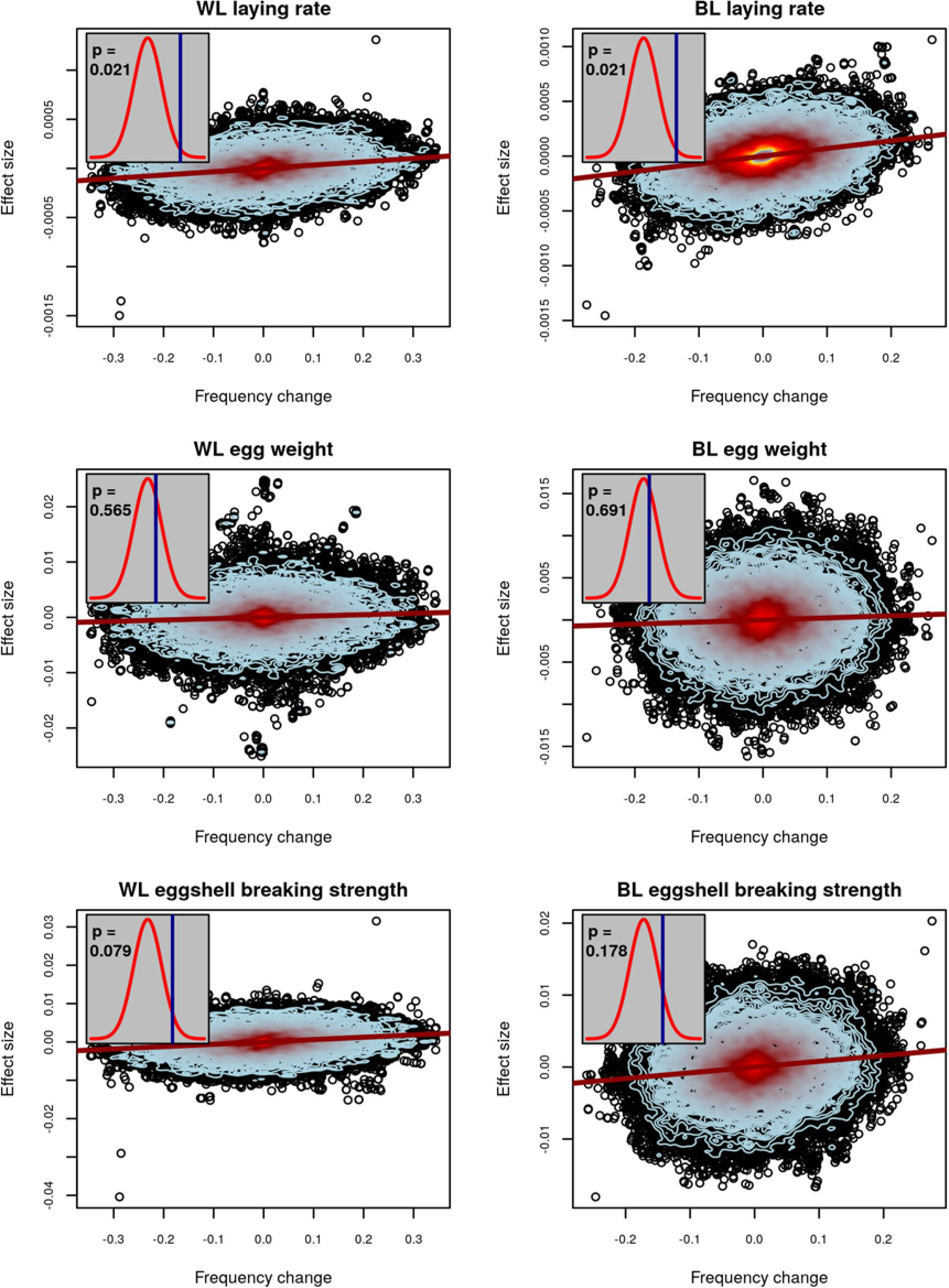
Evidence of selection for chicken traits. For three traits in white (left column) and brown (right column) laying hens, the relationship between estimated allelic effects at individual SNPs and the change in allele frequency over generations is plotted. The red line is a regression of effect size on allele frequency change. Contour lines indicate the density of points, with blue contours indicating fewer points than red. Inset plots depict observed values of Ĝ (blue lines) and their statistical significance based on a comparison to permuted null distributions (red densities) for no-selection scenarios. An exact two-sided p-value is given within each inset. Significant values of Ĝ above the permuted mean indicate selection operated in the positive direction, while significant values below the permutation mean indicated selection operated in the negative direction.

## Discussion

We have defined a test statistic, *Ĝ,* that combines phenotypic and genotypic information to test for selection on traits controlled by many loci of small effect. The approach utilizes estimated effect sizes for individual loci and allele frequency changes across two time-points reflecting possible selection on those loci. Therefore, *Ĝ* is most applicable in experimental or breeding populations, where both pieces of information are readily available via genotyping individuals from multiple generations. However, phenotypic information for estimating allelic effects is only required from a single time-point, so this approach can be applied post-hoc using DNA samples from previous generations even if phenotyping is no longer possible. As the practice of sequencing ancient DNA from archeological sites, museum samples, or other sources becomes progressively commonplace (Orlando *et al.* 2015), it will be interesting to explore whether or not this approach may prove applicable for ecological questions, evolutionary studies, and for human research. However, simulations showed a decrease in power as the number of post-selection generations increased, so there is a limit to how far back our test statistic can be fruitfully applied.

### Powerful for highly quantitative traits

Methods for mapping genes associated with important traits or for identifying loci that are under selection are most powerful for large-effect genes. A simple explanation for the disappointing number of associations that have been uncovered to date through GWAS is that complex traits are often controlled by many genes of small effect (Yang *et al.* 2011). If this is the case, enormous sample sizes are required to map loci regardless of the methodological enhancements that can be applied. Human geneticists have had success studying complex traits by utilizing extremely large sample sizes (Rietveld *et al.* 2013; Wood *et al.* 2014). But, sample sizes of this magnitude are not yet achievable within resource limitations for most species, and, arguably, will never be. Conversely, population genetic studies aiming to scan for selection have been most successful at identifying hard sweeps, where a new mutation of large effect rapidly rises to fixation as a result of selection (Pritchard *et al.* 2010). Only few methodologies with limited power exist for mapping soft sweeps, when the beneficial allele is already at intermediate frequency at the start of selection (Garud *et al.* 2015); Ma *et al.* 2015). A likely explanation for the presence of soft sweeps is that they often result from loci of small effect increasing in frequency slowly in a population and therefore existing on multiple distinct haplotypes or mutating multiple times before fixation. In an agricultural context, many soft sweeps may be due to newly defined breeding goals which put selection pressure on genes that previously were segregating in the populations, but were selectively neutral. The *Ĝ* statistic does not attempt to map specific genes—instead it pools information from all SNPs to test for selection on specific traits. This approach completely avoids the question of which loci are associated with a trait. Instead of testing each SNP, we perform one test based on information from all SNPs. Therefore, a strong statistical signal arises when a large proportion of SNPs behave similarly but not when a few SNPs portray strong signals on their own. That said, researchers are often interested in identifying selected traits whether they correspond to selection on many genes at once or simply a few large-effect genes. In this case, the implementation of our *Ĝ* test in conjunction with a traditional selection-mapping approach aimed at identifying selected loci will likely together be powerful for identifying selection regardless of the underlying genetic architecture (Figure 1).

It was recently argued that most complex disease traits in humans are controlled by small-effect genes dispersed throughout the genome (Boyle *et al.* 2017). Likewise, many important traits in agricultural animal and plant species tend to be quantitative in nature and are presumably controlled by small-effect genes (Goddard and Hayes 2009; Wallace *et al.* 2014). For these agricultural organisms, geneticists and breeders have long recognized the benefits that can be achieved by predicting breeding values and/or phenotypes based on models that use all SNPs simultaneously (Meuwissen *et al.* 2001; Heffner *et al.* 2009; Goddard and Hayes 2009). In fact, the development of these models has led to dramatic re-designs of modern breeding protocols (Schaeffer 2006; Cabrera-Bosquet *et al.* 2012). The *Ĝ* statistic represents one avenue to leverage information from all measured SNPs to gain an understanding of the evolutionary history of a population. This approach is analogous to genomic selection/prediction as utilized by animal and plant breeders, with an important distinction: instead of predicting breeding values to determine which individuals should be selected for the future, it utilizes genotypic frequencies over time coupled with phenotypic information to unravel the history of selection in the past.

### Genotypes from base population provide high power

Compared to other methods that test for selection on quantitative traits (Berg and Coop 2014; Zeng *et al.* 2017), *Ĝ* leverages genotypic information from multiple time points and that it incorporates information from all SNPs instead of restricting to a previously identified set of SNPs from one or multiple independent GWAS’s. With the exception of a few traits in heavily studied species, such as human height (Wood *et al.* 2014), few species, if any, provide the enormous sample sizes required to implicate a large number of loci for any quantitative traits. This includes situations where scientists are reasonably certain that a genetic architecture consisting of small-effect loci persists. Importantly, *Ĝ* is powerful because of the independence of the estimation of allele frequency changes across generations and effect sizes, respectively. Even when allelic effects and/or allele frequency changes are small, they cumulatively generate a powerful test since they can be compared across all genotyped loci. However, our analysis of the chicken data suggested that the power of the test can be reduced through noisy estimation of allele frequency change. Our reliance on pedigree data to derive initial allele frequencies was not as precise as the direct measurement of initial allele frequencies that was conducted for maize. Although we were still able to find evidence of selection on traits including laying rate, which was almost certainly under the strongest selection, there were selected traits we did not detect potentially due to this noise.

### Future directions and conclusions

The use of *Ĝ* to test for selected traits avoids the requirement of preliminarily identifying candidate genes or regions. Therefore, the approach is particularly applicable in experimental, agricultural, and natural populations for which available resources dictate limited sample sizes for conducting massive mapping studies for such preliminary identification. In contrast to purely population-genetic analyses, which rely solely on genotypic information, the method requires that phenotypic data be collected from at least one time-point of genotyped individuals. Additionally, two time-points of genotypic information are needed, either directly or through pedigree-based imputation.

While the *Ĝ* statistic is most directly applicable for the discovery of traits that have been previously under selection during recent evolution, it may have additional applications. Recent studies have demonstrated that distinct physical regions of the genome, such as individual chromosomes, often contribute a disproportionate amount to trait variance (Bernardo and Thompson 2016). Rather than applying the *Ĝ* statistic genome-wide, future research should be done to determine whether it can be applied across any collections of loci such as individual chromosomes, pathways, gene-families, functional classes, or other categories to test if these show evidence of selection on a quantitative trait. This would represent a process allowing researchers to map significant features as opposed to individual genes. Likewise, thus far we have estimated the direction of selection (positive or negative) from *Ĝ*, but not the magnitude. Further research should be performed to determine whether or not this or a similar statistic can be used to recapitulate the selection gradient.

As it stands, using *Ĝ* simply to identify traits that have been under selection in the past may prove enormously useful. Whether agricultural, experimental, or natural, it is often difficult to determine all of the traits that are advantageous in a population or respond to natural or anthropogenic selection, including undesired selection responses. The application of the *Ĝ* statistic genome-wide allows this determination, which may help scientists select the right traits for maximum agricultural production, determine inadvertently selected lab traits impacting experimental outcomes, and establish ecologically important traits for survival in the wild.

## Materials and methods

### Simulations

Each simulation started with a random mating historical population. After 5 thousand generations, selection began and simulations proceeded with more control over each generation. Truncation selection was performed based on high phenotype. Drift simulations were identical to selection simulations in terms of genome layout and genetic basis of the trait, but individuals were selected randomly. Simulations were performed with QMSim (Sargolzaei and Schenkel 2009). Parameters for our generic simulation model are provided in full in Supplemental Table 1. We varied specific parameters as follows:

#### Number of QTL

Genetic architectures with 10, 50, 100, 1,000, or 10,000 QTL were simulated.

#### Number of individuals phenotyped

After selection was simulated, the phenotypes from a subset included 1,000, 500, 250, 100, or 50 individuals were sampled and used for estimating SNP effects.

#### Selection intensity

The number of males and females reproducing each generation was always simulated to be 500, respectively. To vary selection intensity, we simulated litter sizes of 4, 20, 40, and 200.

#### Number of generations of selection

Selection simulations were conducted for 1, 10, 20, 50, and 100 generations.

#### Phenotyping generation

For 20-generation simulations, phenotypes were analyzed from pre-selection individuals (generation 0), mid-selection individuals(generation 10), and post-selection individuals (generation 20).

#### Number of generations after selection

After 20 generations of selection, we evaluated whether *Ĝ* was still significant after 5, 20, 50, or 100 generations without selection.

### Selection mapping in simulations

For the set of simulations where number of QTL were varied, pre- and post-selection simulated allele frequencies were output from QMSim. These were used to calculate marker-specific *F_ST_* values, as was performed by (Lorenz *et al.* 2015). *F_ST_* was computed according to 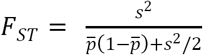, where *s^2^* is the sample variance of allele frequency between pre- and post-selection populations, and 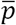 is the mean allele frequency (Weir and Cockerham 1984). Experiment-wide 5% significance threshold were identified based on the 95% *F_ST_* quantile observed from drift simulations. These thresholds were applied to *F_ST_* values obtained from selection simulations to determine detection and false positive rates. Simulated QTL were declared detected if a significant marker was identified within a .1 cM window surrounding the QTL. False positives were defined as markers that were not within a .1 cM window surrounding any simulated QTL.

### Maize data

All maize data were previously published and described by Lorenz et al. (2015). In brief, a selection index comprised of silage-quality traits was used to perform reciprocal recurrent selection. Traits comprising the index were yield, dry matter content, neutral detergent fiber (NDF), protein content, starch content, and *in-vitro* digestibility (www.cornbreeding.wisc.edu). Phenotypic data included five cycles of selection, encompassing approximately 20 generations in total. Tens to hundreds of individuals were sampled from each cycle of selection to be genotyped. Genotyping was performed with the MaizeSNP50 BeadChip, which includes 56,110 markers in total (Ganal *et al.* 2011). After removing monomorphic SNPs, redundant SNPs, quality filtering, and imputing, as described in Lorenz (2015), 10,023 informative SNPs remained.

Allele frequencies were computed for each cycle of selection. Because only 5 and 11 individuals from cycles 0 and 1 were genotyped, respectively, allele frequency change from cycle 2 (n = 163) to cycle 5(n = 211) was computed for each SNP. Since all SNPs were di-allelic, the frequency of only one allele was tracked, and the frequency change for that allele perfectly mirrored the change for the other allele. For the tracked allele only, allelic effects were estimated using the R package RR-BLUP (Endelman 2011). Phenotypic information was available from individuals representing selection cycles 1 through 4, and since population size was small we used all phenotyped individuals to estimate SNP effects. To accomplish this without biasing effect estimates due to drift, a fixed effect for cycle was included in our model. Our exact analysis scripts are available at github.com/timbeissinger/ComplexSelection.

### Chicken data

Data were available for one white layer (WL) and one brown layer (BL) line from a commercial breeding program. Both closed lines have been selected over decades with a similar composite breeding goal, comprising, among others, laying rate, body weight and feed efficiency of the hens, as well as egg weight and egg quality, where the respective weights of the different traits varied between lines and over time. In total, 673 (743) WL (BL) individuals were genotyped, of which > 80% were from the last generation and the remaining animals were parents, grand-parents, and great-grandparents of the actual birds. For all genotyped individuals, complete pedigree data were available comprising 2109 (1879) individuals and going 13 (9) generations back in WL (BL). The oldest generation was defined as the base population and comprised 111 (64) ungenotyped individuals being separated from the majority of genotyped individuals by 12 (8) generations.

Current individuals were genotyped with the Affymetrix Axiom® Chicken Genotyping Array which initially carries 580K SNPs. This data were pruned by discarding sex chromosomes, unmapped linkage groups, and SNPs with minor allele frequency (MAF) lower than 0.5% or genotyping call rate smaller than 97%. Individuals with call rates smaller than 95% were also discarded. Subsequently, missing genotypes at the remaining loci were imputed with Beagle version 3.3.2 (Browning and Browning 2009),resulting in sets of 277,522 (334,143) SNPs for the WL (BL) individuals.

To calculate the allele frequency change in the chicken populations, the allele frequency in the base population individuals had to be reconstructed by statistical means. This was done with the approach of Gengler *et al.* (2007), which, in short, considers the allele frequency in an individual as a quantitative and heritable trait and uses a mixed model approach to obtain a best linear unbiased prediction (BLUP) for the allele frequency of all un-genotyped individuals. This is done by linking the genotyped offspring to the un-genotyped ancestors via the pedigree information (for details, see Gengler *et al.* 2007). This required solving 277,522 (334,143) linear equation systems of dimension 2109 (1879) for the WL (BL) data set. Next, *Δ_i_* for locus *i* was calculated as the difference of the observed allele frequency of the genotyped individuals in the current and the 3 ancestral generations and the average estimated allele frequency of the 111 (64) base population individuals 12 (8) generations back.

For each genotyped individual, conventional (non-genomic) BLUP breeding values and the respective reliabilities for a wide set of traits were available. SNP effects were estimated in a two-step procedure: first, for each trait in each line genomic breeding values were estimated via genomic BLUP (GBLUP), followed by a back-solution of estimated SNP effects. In the GBLUP step, the model ***y*** = **1***μ* + ***Zg*** + ***e***, was solved, where ***y*** is the vector of de-regressed proofs [DRPs] of genotyped individuals for a specific trait; *μ* is the overall mean; ***g*** is the vector of additive genetic values (i.e. genomic breeding values) for all genotyped chickens; ***e*** is the vector of residual terms; **1** is a vector of 1s and ***Z*** is a squared design matrix assigning DRPs to additive genetic values with dimension number of all genotyped individuals. Residual terms were assumed to be distributed 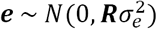, where ***R*** is a diagonal matrix with diagonal elements 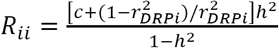(Garrick *et al.* 2009) for an individual *i* in the training set, where 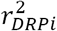 is the reliability of DRP for individual *i*, 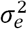 is the residual variance, using *c* set to 0.1. The distribution of additive genetic values is assumed to be 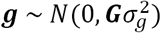, where 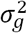 is the additive genetic variance and *G* is a realized genomic relationship matrix which was constructed according to (VanRaden 2008). Estimation of variance components and genomic breeding values was done with ASReml 3.0 (Gilmour et al., 2009).

Next, estimated SNP effects *ŝ* were obtained following Strandén and Garrick (2009) as

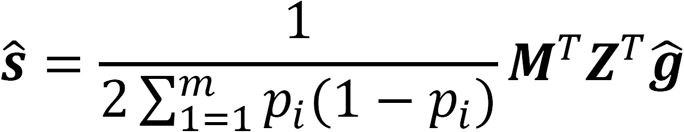

where ***M*** is a matrix of dimension number of genotyped individuals x number of genotyped SNPs with entry *m_ij_ = x_ij_ — 2p_j_* where *x_ij_* is the genotype of individual *i* at locus *j* (coded as 0, 1, or 2 which are counts of the reference allele) and *p_j_* is the population frequency of the reference allele at SNP *j.*

## Computational Resources

Computation was performed using the University of Missouri Informatics Core Research Facility BioCluster (https://bioinfo.ircf.missouri.edu/). Computational nodes where simulations were performed had 64 cores and 512 GB of RAM. Analysis of maize and chicken data was performed on a mediocre laptop with 8 GB of RAM.

## Data availability

Maize data are available in from Lorenz et al. (2015). Chicken data, including allele frequency change and estimated SNP effects, are available at Figshare with DOI 10.6084/m9.figshare.5899267. All scripts used for simulations and analysis are available at github.com/timbeissinger/ComplexSelection.

## Acknowledgements

We thank Natalia de Leon, Aaron Lorenz, and Lohmann GMBH for generating the maize and chicken biological data used in this study. We are grateful for helpful discussions with Emily Josephs and Aaron Lorenz. This research was supported by the USDA Agricultural Research Service, CRIS project number 5070-21000-038-00-D.

---

**Supplemental Table 1:** Simulation parameters.

| Parameter |  | Value(s) |
| --- | --- | --- |
| **Genetic basis of the trait** |  |  |
| Heritability |  | 0.5 |
| QTL Heritability |  | 0.5 (all heritability attributable to QTL) |
| Phenotypic variance |  | 1 |
| **Historical Population** |  |  |
| Population size |  | 10,000 |
| Number of generations |  | 5,000 |
| Marker mutation rate (only historical gens) |  | 2.5e-5 |
| QTL mutation rate (only historical gens) |  | 2.5e-5 |
| **Breeding (selected) Population** |  |  |
| Number of selected males/generation |  | 500 |
| Number of selected females/generation |  | 500 |
| Litter size |  | 10 |
| Number of generations |  | 20 |
| Mating design |  | Random union of gametes, discrete generations |
| **Genome** |  |  |
| Number of chromosomes |  | 10 |
| Chromosome size |  | 100 cM |
| Markers/chromosome |  | 10,000 |
| Marker spacing |  | Even |
| Alleles/marker |  | 2 |
| Marker allele frequencies |  | Random (uniformly distributed) |
| Number of QTL |  | 1,000 |
| QTL spacing |  | Even |
| Alleles/QTL |  | 2 |
| QTL allele frequencies (in first gen) |  | Equal (0.5) |
| QTL allele effects |  | Random (uniformly distributed) |

**Supplemental Figure 1:**
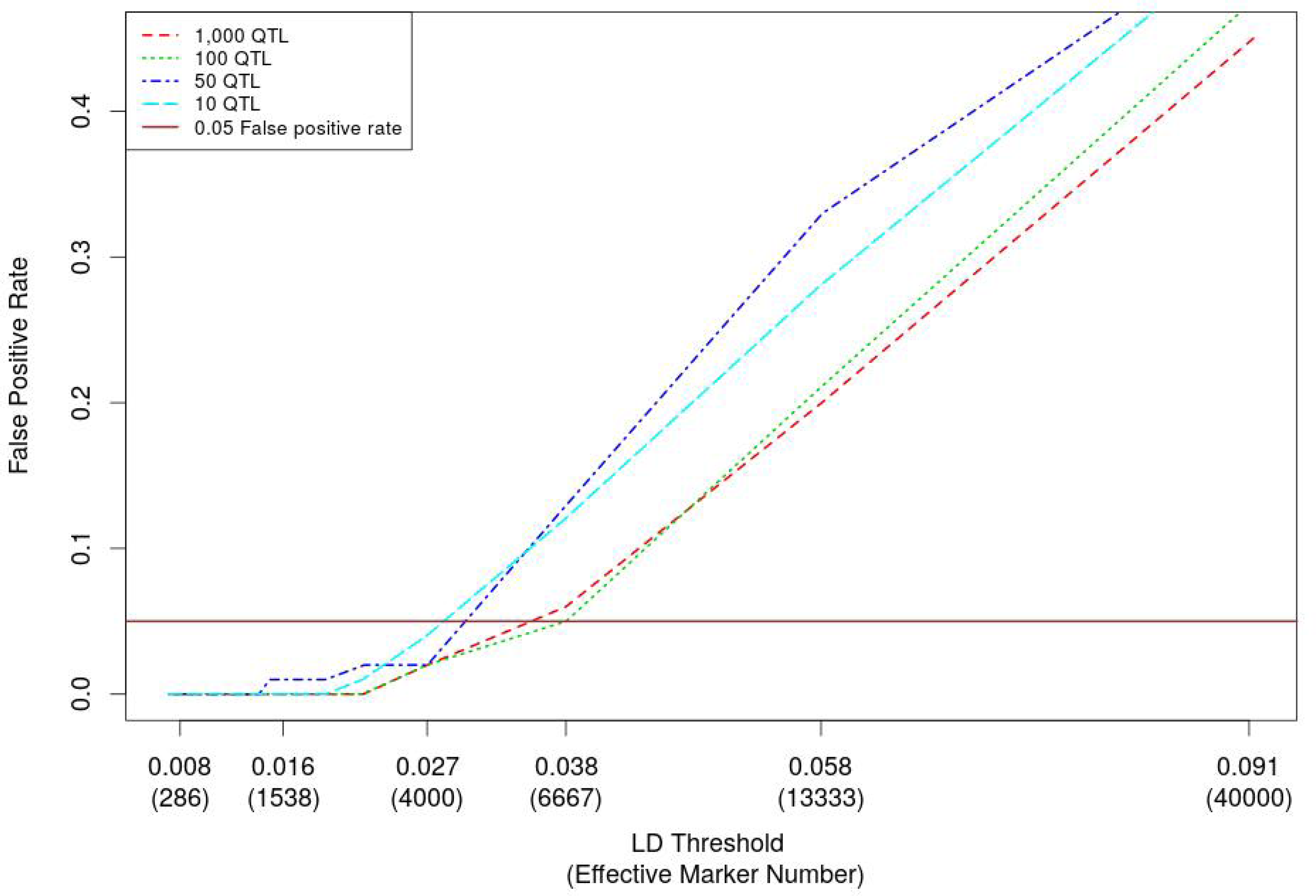
False positive rate depends on the *number of effective markers.* The y-axis of this plot shows the false positive rate for simulations of different genetic architectures that was realized with varying effective numbers of markers. The x-axis depicts the mean LD-threshold across simulations that corresponded to a particular effective number of markers. Simulations suggested that defining the effective number of markers as the number of genome-segments such that LD across each segment is expected to be in the interval *R^2^* ∈ [0.027, 0.038] appropriately controls false positive rate.

**Supplemental Figure 2:**
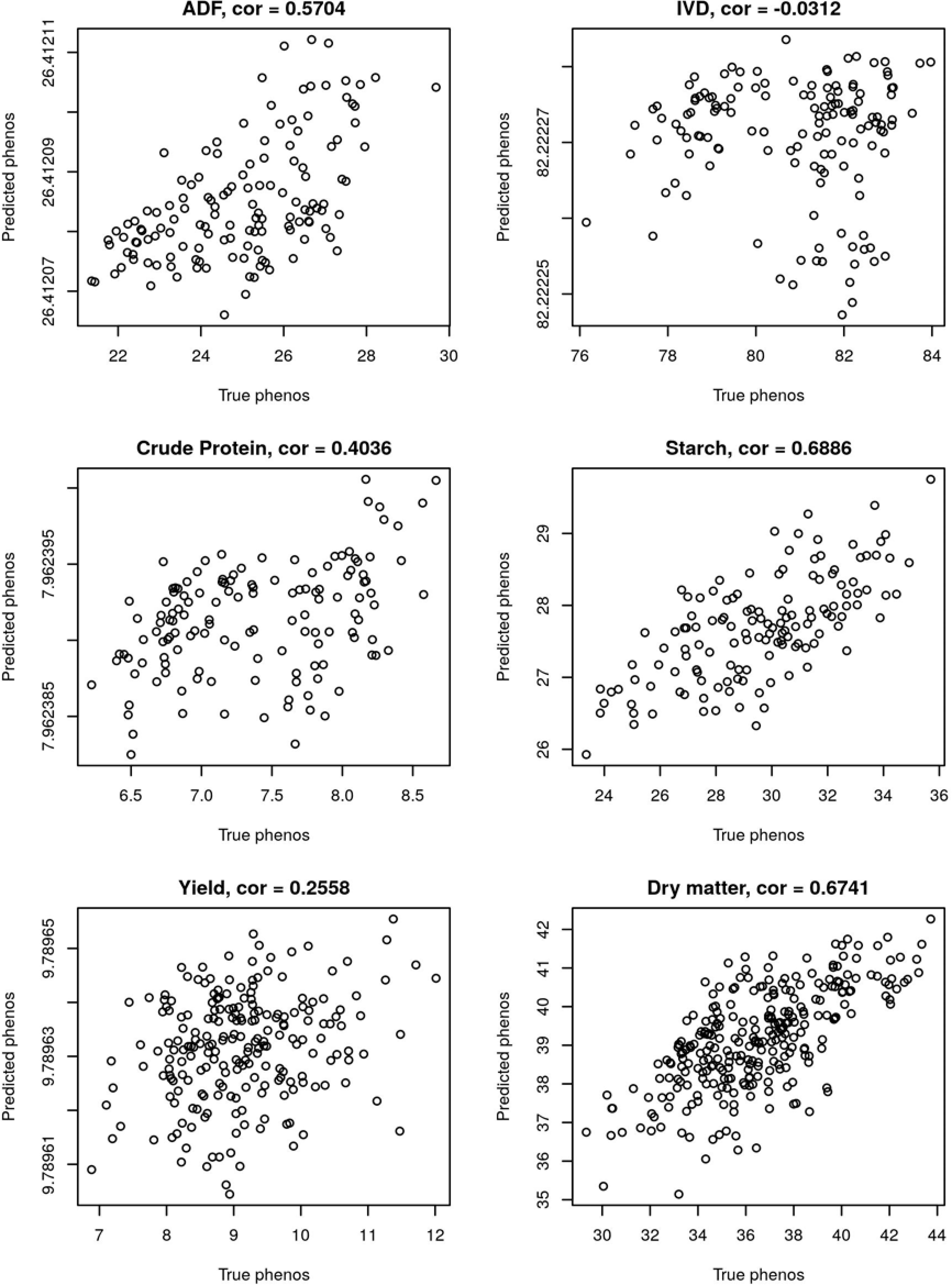
Correlation between predicted and observed phenotypes when RRBLUP was used for genomic prediction in the maize dataset.

**Supplemental Figure 3:**
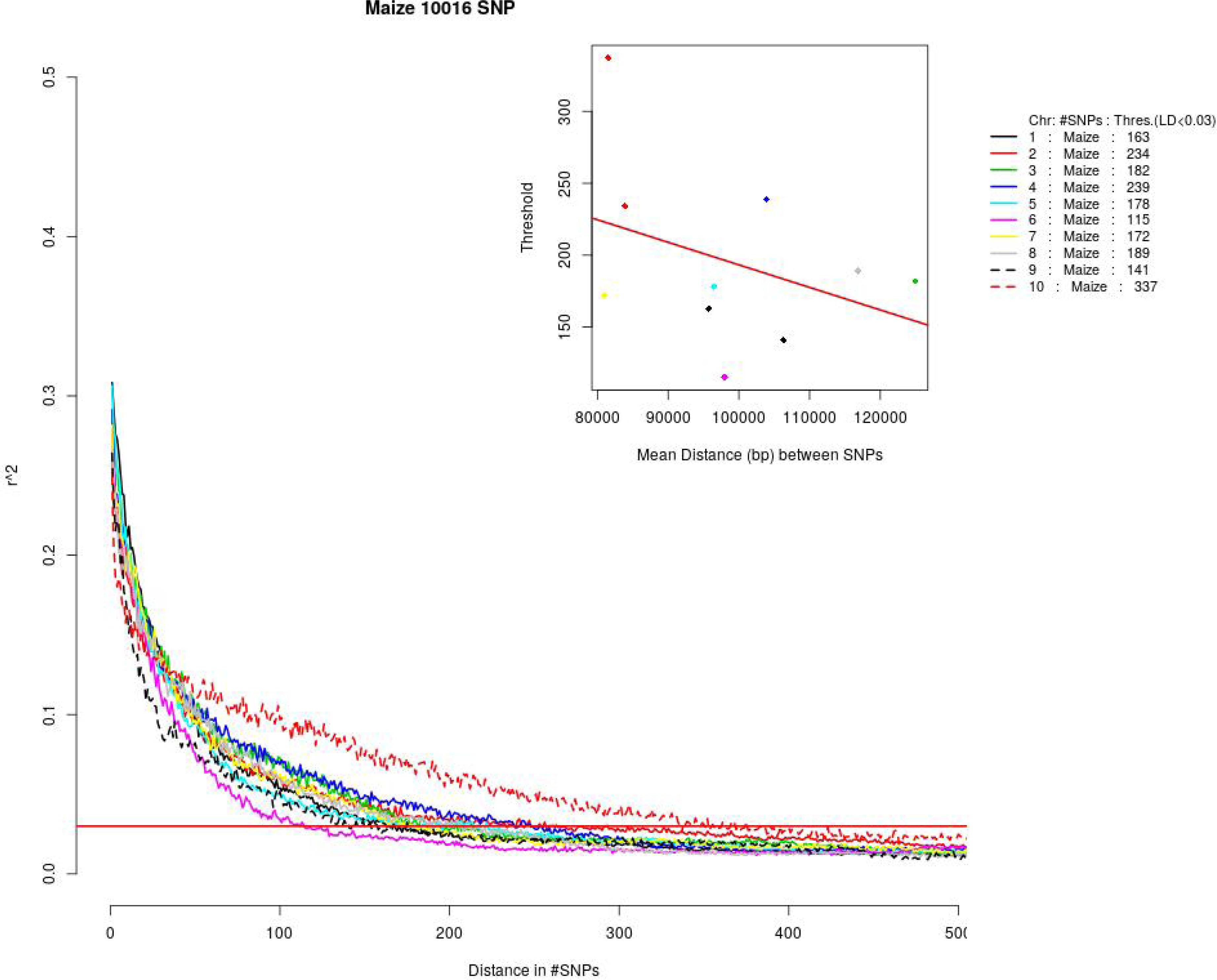
LD Decay by chromosome in the WQS maize population. For each chromosome, LD is plotted against the distance between SNPs (in number of markers). The effective number of independent markers (*m_ind_*) for our test was determined by dividing the total number of markers by the mean distance between markers such that R^2^ ≤ 0.03.

**Supplemental Figure 4:**
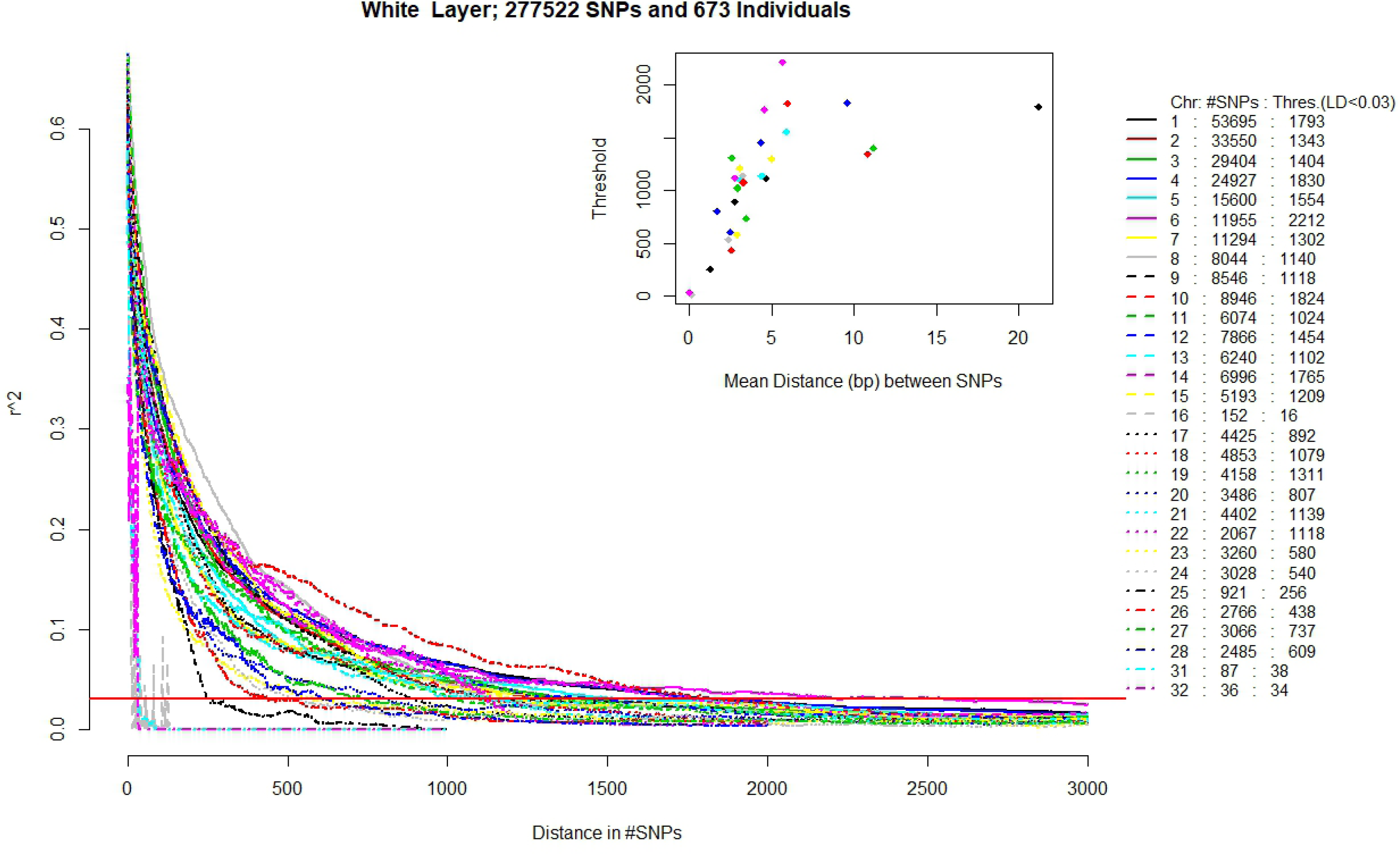
LD Decay by chromosome in the White Layer chicken population. For each chromosome, LD is plotted against the distance between SNPs (in number of markers). The effective number of independent markers *(m_ind_)* for our test was determined by dividing the total number of markers by the mean distance between markers such that R^2^ ≤ 0.03.

**Supplemental Figure 5:**
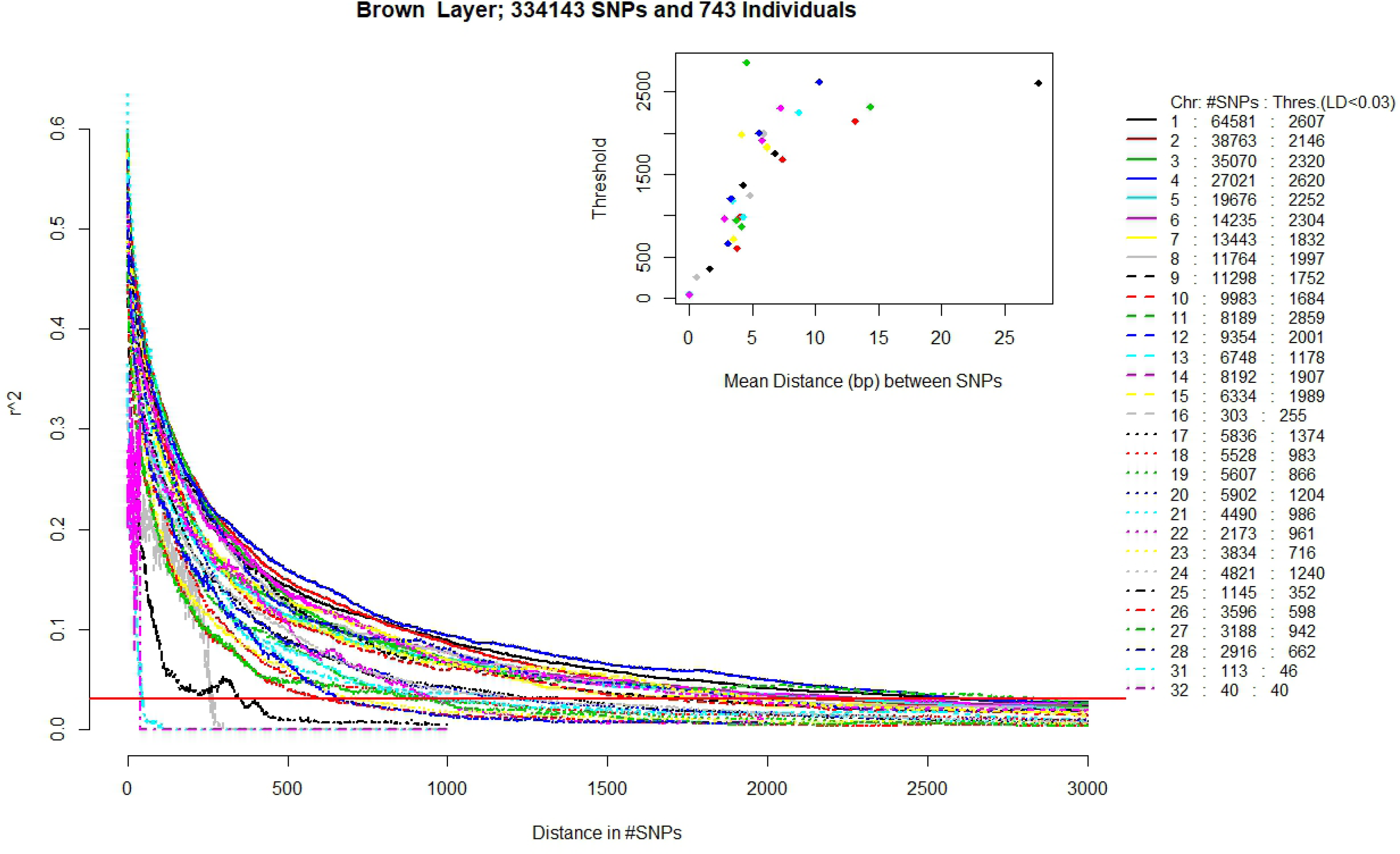
LD Decay by chromosome in the Brown Layer chicken population. For each chromosome, LD is plotted against the distance between SNPs (in number of markers). The effective number of independent markers *(m_ind_)* for our test was determined by dividing the total number of markers by the mean distance between markers such that R^2^ ≤ 0.03.

**Supplemental Figure 6:**
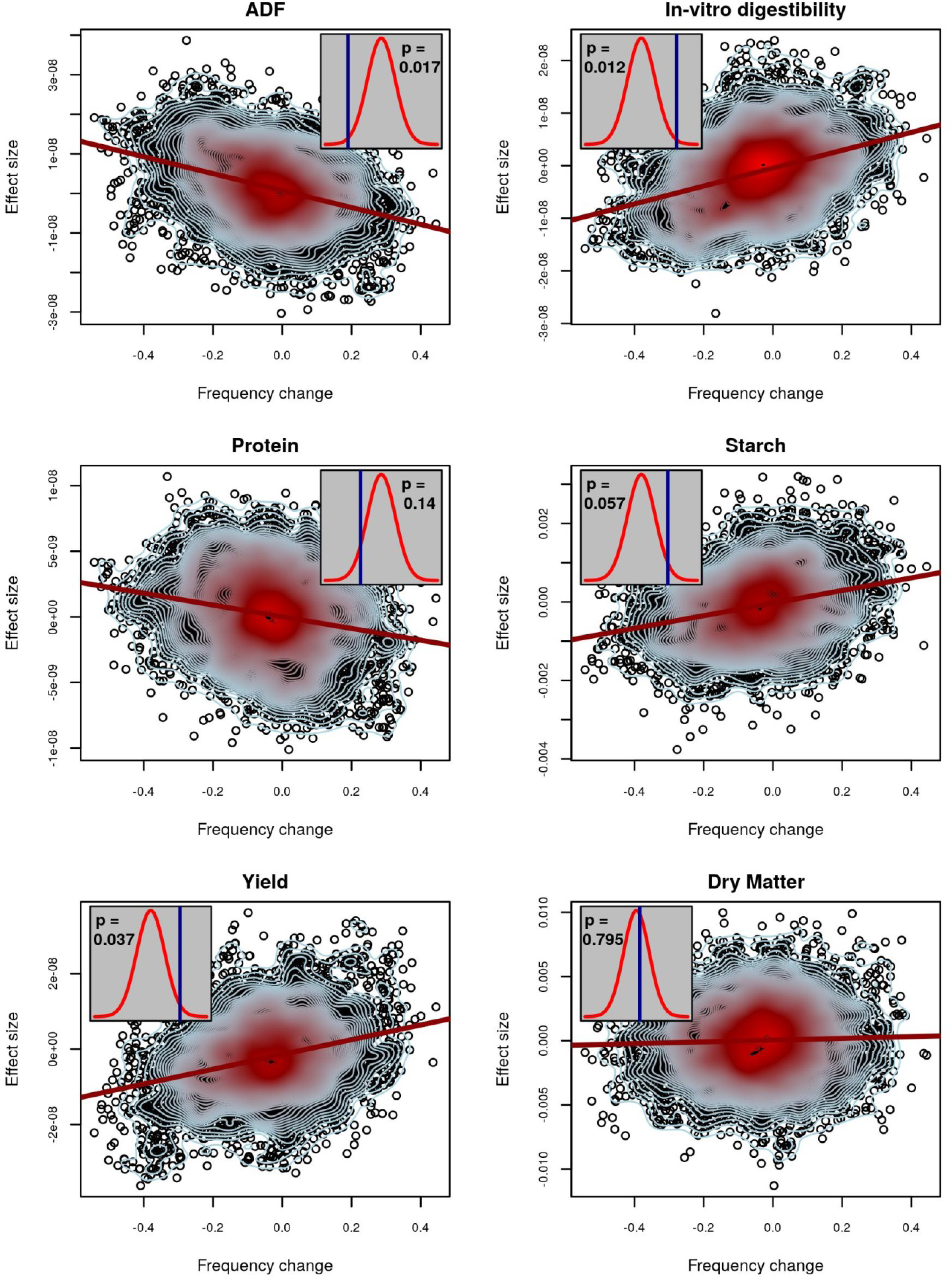
Evidence of selection for maize silage traits. SNPs with minor allele frequency <0.05 were removed for this analysis. For six traits, the relationship between estimated allelic effects at individual SNPs and the change in allele frequency over generations is plotted. The red line is a regression of effect size on allele frequency change. Contour lines indicate the density of points, with blue contours indicating fewer points than red. Inset plots depict observed values of Ĝ (blue lines) and their statistical significance based on a comparison to permuted null distributions (red densities) for no-selection scenarios. An exact two-sided p-value is given within each inset. Significant values of G above the permuted mean indicate selection operated in the positive direction, while significant values below the permutation mean indicated selection operated in the negative direction.

**Supplemental Figure 7:**
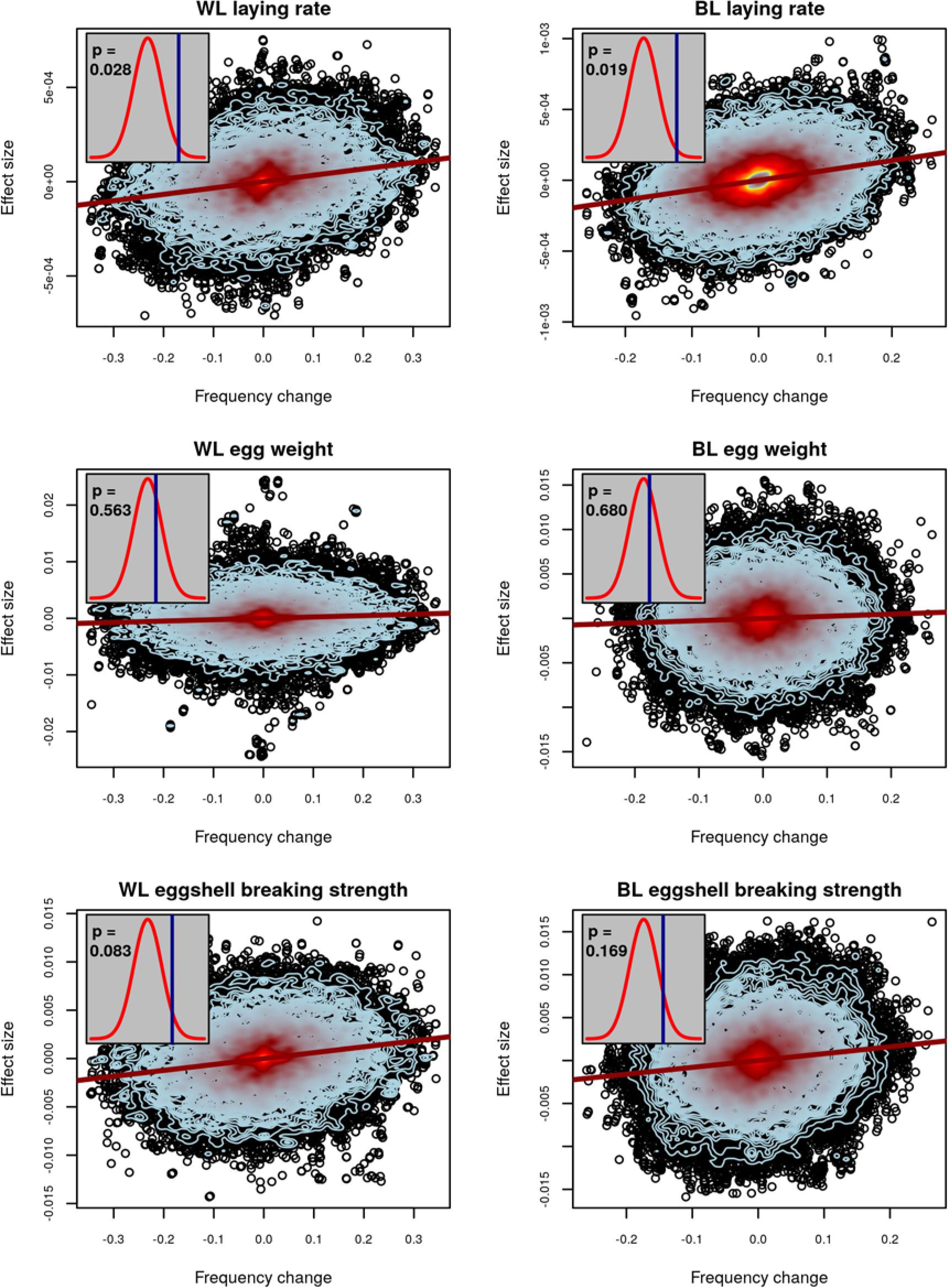
Evidence of selection for chicken traits, with potential outliers removed. This plot demonstrates a reanalysis of the chicken data shown in Figure 3 after removing of the 10 SNPs with the largest-magnitude effect size for each trait. For three traits in white (left column) and brown (right column) laying hens, the relationship between estimated allelic effects at individual SNPs and the change in allele frequency over generations is plotted. Contour lines indicate the density of points, with blue contours indicating fewer points than red. Inset plots depict observed values of Ĝ (blue lines) and their statistical significance based on a comparison to permuted null distributions (red densities) for no-selection scenarios. An exact two-sided p-value is given within each inset. Significant values of Ĝ above the permuted mean indicate selection operated in the positive direction, while significant values below the permutation mean indicated selection operated in the negative direction.

